# Colonization history of the Western Corn Rootworm (*Diabrotica virgifera virgifera*) in North America: insights from random forest ABC using microsatellite data

**DOI:** 10.1101/117424

**Authors:** Eric Lombaert, Marc Ciosi, Nicholas J. Miller, Thomas W. Sappington, Aurélie Blin, Thomas Guillemaud

## Abstract

First described from western Kansas, USA, the western corn rootworm, *Diabrotica virgifera virgifera*, is one of the worst pests of maize. The species is generally thought to be of Mexican origin and to have incidentally followed the expansion of maize cultivation into North America thousands of years ago. However, this hypothesis has never been investigated formally. In this study, the genetic variability of samples collected throughout North America was analysed at 13 microsatellite marker loci to explore precisely the population genetic structure and colonization history of *D. v. virgifera*. In particular, we used up-to-date Approximate Bayesian Computation methods based on random forest algorithms to test a Mexican versus a central-USA origin of the species, and to compare various possible timings of colonization. This analysis provided strong evidence that the origin of *D. v. virgifera* was southern (Mexico, or even further south). Surprisingly, we also found that the expansion of the species north of its origin was recent - probably not before 1100 years ago - thus indicating it was not directly associated with the early history of maize expansion out of Mexico, a far more ancient event.

## Introduction

The western corn rootworm (WCR), *Diabrotica virgifera virgifera,* is a major economic pest of maize, *Zea mays,* in North America and, since the end of the twentieth century, in Europe (Gray et al. 2009; Vilà et al. 2009). Although the invasion history of WCR in Europe has been well investigated (Miller et al. 2005; Ciosi et al. 2008), its biogeography, colonisation history and potential association with maize domestication in America are poorly understood.

Because of the geographical distribution of most other diabroticites and the close association of WCR with maize, the species is commonly considered as originating from Mexico, or possibly Guatemala, where its original native host was probably *Tripsacum,* a close wild relative of maize (Smith 1966; Branson and Krysan 1981; Gray et al. 2009). The classically proposed scenario is that WCR fed on early domesticated maize, and incidentally followed the dissemination of the plant into southwestern North America and the Great Plains, so that the history of WCR tracks the history of maize into those regions (Branson and Krysan 1981). Maize is a human-made variant of teosinte which was domesticated about 9,000 years before present (BP) in southern Mexico (Matsuoka et al. 2002; Buckler and Stevens 2005). The cultivation of maize slowly expanded northward to reach the present-day states of Arizona and New Mexico, USA around 4,100 BP (Merrill et al. 2009; da Fonseca et al. 2015), and became an important part of the diet of some groups in the Four Corners region between 2,400 and 3,000 BP (Coltrain et al. 2010; Smith 2017). The selection of new variants that were better adapted to temperate climates helped to spread maize further into the northern USA and Canada by around 2,000 years BP (Fritz 1990; Hart et al. 2007; Tenaillon and Charcosset 2011), but it was a minor crop throughout America north of Mexico before 900 to 1000 CE (Boyd et al. 2008; Simon 2017; Smith 2017). A large increase in maize cultivation by European migrants in North America occurred in the nineteenth century, probably helped by development of new cultivars (Anderson and Brown 1952; Doebley et al. 1988). Finally, the intensification of cultivation in the mid-20th century coinciding with commercialization of modern inbred hybrids widely boosted this trend (Kutka 2011).

However, different WCR origin scenarios are possible, such as a far more recent colonization history than that of maize, and/or a more northern North American origin of the species. These scenarios are based on the dates of first observation of WCR in America and on the knowledge of its ecology. *D. virgifera* was first described by Le Conte from two individuals collected in 1867 from blossoms of *Cucurbita foetidissima* in western Kansas (Le Conte 1868; Metcalf 1983; Krysan and Smith 1987), and the first economic damage on maize was noticed only in 1909 in Colorado (Gillette 1912). The species is known to have been present in more southern States such as Arizona and New Mexico, as well as in Mexico, at least since the end of the nineteenth century (Horn 1893), but more detailed information about their presence in these areas is not available before the 1950s (Chiang 1973; Krysan and Smith 1987). The colonization of the Eastern USA and Canada by WCR has been well monitored and is very recent compared to the widespread cultivation of maize in those areas beginning around 1000 CE: beginning in the 1940s, WCR started to spread eastward from the western Great Plains at considerable speed to reach the East coast of North America in the mid-1980s (Krysan and Smith 1987; Gray et al. 2009; Meinke et al. 2009). Furthermore, behavioural data do not fully support an exclusive shared history between WCR and maize, suggesting instead a host switch, which could possibly be recent, from a very different host plant (than *Tripsacum)* to maize, either in Mexico or the central USA. Indeed, larvae have no mechanism for distinguishing maize from a distance (Branson and Krysan 1981), whereas WCR adults are strongly attracted to cucurbitacins, secondary metabolites of Cucurbitaceae (Metcalf and Lampman 1989). Potential alternative hosts in North America include a number of native grass species (Clark and Hibbard 2004; Oyediran et al. 2004), but their current importance in a maize-dominated agroecosystem is probably minimal (Moeser and Hibbard 2005; Campbell and Meinke 2006).

In this study, we characterized the current genetic structure of WCR in North America, from Mexico to the northeastern USA, by Bayesian clustering methods and more classical population genetic statistics and methods. We then performed up-to-date random forest approximate Bayesian computation analyses to quantitatively compare colonization scenarios of WCR populations in North America.

## Methods

### Sampling, genotyping and genetic variation

Nine hundred and seventeen WCR adults were collected from 21 sites (14 to 62 WCR per site) in North America between 1998 and 2006, covering a substantial part of the distribution of this species in America (Fig. 1; Table S1). Samples from twelve of these sites were used in previous studies (Table S1; Kim and Sappington 2005; Kim et al. 2008; Coates et al. 2009). Genotyping at 13 microsatellite marker loci was carried out in three separate multiplex PCRs for all individuals as described by Bermond *et al.* (2012).

**Figure 1:**
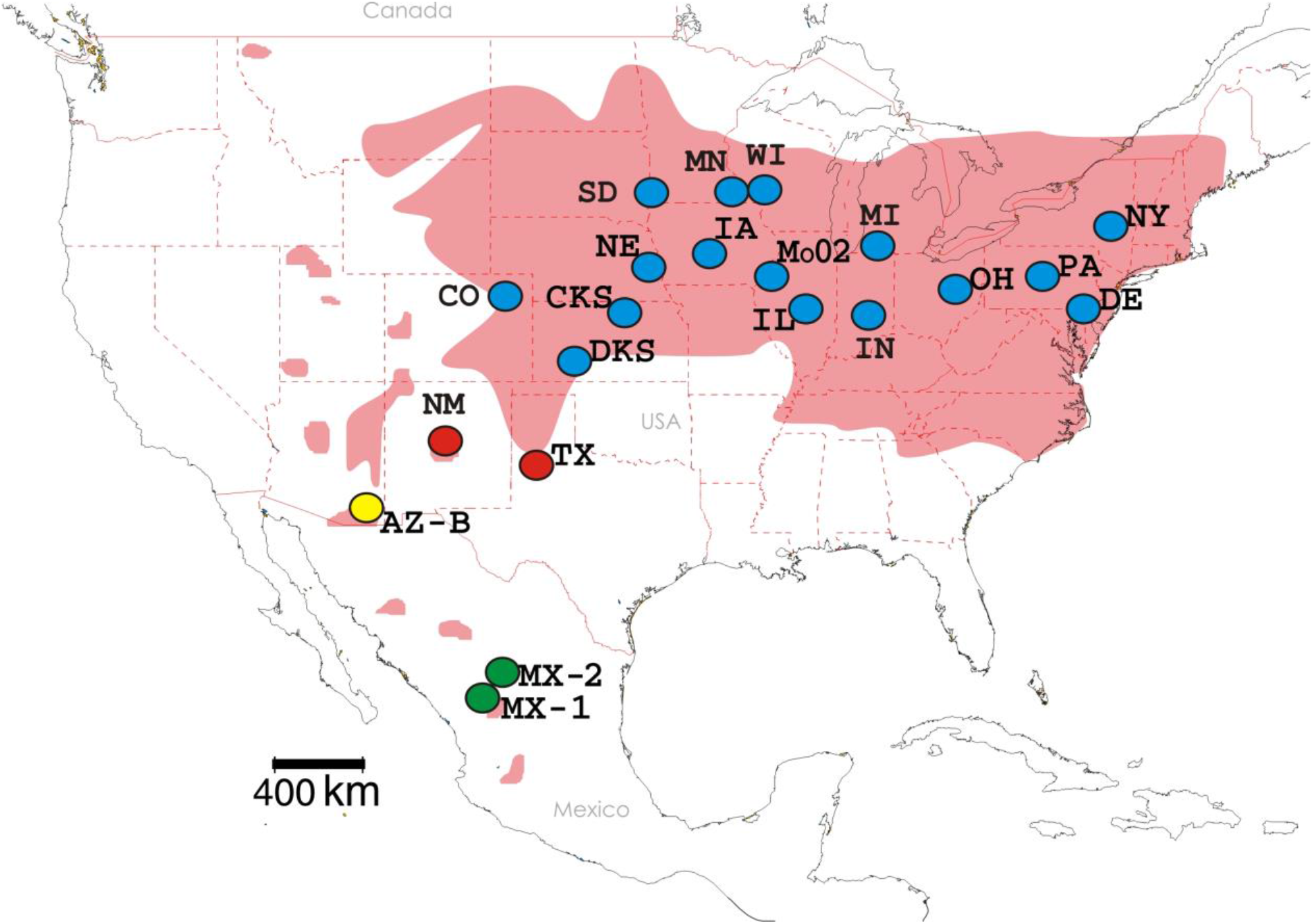
Geographic locations of genotyped site-samples of WCR and genetic units inferred from Bayesian clustering analyses.

Notes: Site-sample names are as in Table S1. The pink areas roughly correspond to the geographic distribution of WCR in North America. Site-samples of the same color belong to the same genetic unit, as assessed by hierarchical procedures applied to the Bayesian clustering methods implemented in STRUCTURE and BAPS (Figures S3 and S4): “Mexico” in green, “Arizona” in yellow, “New Mexico/Texas” in red and “Colorado/New York” in blue.

Genetic variation within and between the 21 site-samples were quantified by calculating the mean number of alleles per locus *NA,* the mean expected heterozygosity *He* (Nei 1987) and pairwise *F*_ST_ estimates (Weir and Cockerham 1984) using Genepop (version 4.2, Raymond and Rousset 1995). To take into account the differences in sample size between site-samples, we computed the mean allelic richness *(AR)* corrected for 10 individuals by the rarefaction method (Petit et al. 1998) with HP-Rare (version 1.1, Kalinowski 2005). Hardy-Weinberg and genotypic differentiation tests were performed using Fisher exact tests implemented in Genepop (version 4.2, Raymond and Rousset 1995), and significance levels were corrected for multiple comparisons biases by the false discovery rate procedure (Benjamini and Hochberg 1995). Null allele frequencies for each locus and each site-sample were estimated following the expectation maximum algorithm of Dempster *et al.* (1977) using FreeNA (Chapuis and Estoup 2007). We constructed a neighbour-joining (NJ) tree (Saitou and Nei 1987) using pairwise genetic distances as described by Cavalli-Sforza and Edwards (1967), using Populations software (version 1.2.30, Langella 1999). The robustness of tree topology was evaluated by carrying out 1,000 bootstrap replicates over loci. Finally, isolation-by-distance was evaluated by determining the correlation between pairwise natural logarithmic geographic distances and genetic distances [*F*_ST_/(1-*F*_ST_)], through a Mantel test with 10,000 permutations implemented in Genepop (version 4.2, Raymond and Rousset 1995).

### Population structure and definition of genetic units

The clustering approach implemented in STRUCTURE (v2.3.4, Pritchard et al. 2000) was used to infer the number of potential genetic units within the North American range of WCR. We chose the admixture model with correlated allele frequencies, and default values for all other parameters of the software. Each run consisted of a burn-in period of 2×10^5^ Markov chain Monte Carlo (MCMC) iterations, followed by 10^6^ MCMC iterations. We carried out 20 replicate runs for each value of the number *(K)* of clusters, with *K* set between 1 and [the number of site-samples considered + 1]. To group each site-sample within its most likely genetic unit, we used the hierarchical approach of Coulon *et al.* (2008) as follows. We first analysed the whole dataset, consisting of 21 site-samples (totalling 917 individuals). If the mean natural logarithm of the likelihood of the data *ln(P(X|K))* was maximal for *K* = 1, then the inferred number of clusters was 1 and we stopped the procedure. Otherwise, we determined the highest level of genetic structure by the ΔK method (Evanno et al. 2005). We then partitioned the previous dataset by assigning each site-sample to the inferred cluster for which the mean individual ancestry was greater than 0.8; site-samples with mean ancestry below 0.8 for all clusters were assigned to a specific “admixed” group. We performed successive independent rounds of STRUCTURE analyses on each subset of the data until *ln(P(X|K))* was maximal for *K* = 1, or until only one site-sample remained.

We also used the clustering approach implemented in BAPS software (v5.2, Corander et al. 2003) as a complement to the STRUCTURE analyses. Although both programs identify population structure by minimizing Hardy-Weinberg and linkage disequilibrium within each of *K* clusters, BAPS uses a fast stochastic-greedy optimisation algorithm instead of the MCMC algorithm used in STRUCTURE (Putman and Carbone 2014). We carried out BAPS analyses on groups of individuals (i.e. site-samples) rather than individuals, with simple model assumptions (i.e. no admixture and uncorrelated allele frequencies). We conducted a series of 20 replicate runs, with the upper limit for the number of clusters set as the actual number of sampled sites. BAPS infers the number of clusters (K is a parameter of the model, unlike in STRUCTURE), but we proceeded to a hierarchical approach as well by performing independent analyses within each inferred cluster until the number of newly inferred clusters was one or until only one site-sample remained.

### ABC-based inferences about colonization history

An approximate Bayesian computation analysis (ABC; Beaumont et al. 2002) was carried out to infer the colonization history of WCR in North America. The populations considered in the ABC analysis corresponded to the genetic units previously identified by the two Bayesian clustering methods (i.e., STRUCTURE and BAPS), and each genetic unit was represented in the analysis by a single site-sample (the “core dataset”, see Results section). ABC is a model-based Bayesian method allowing posterior probabilities of historical scenarios to be computed, based on historical data and massive simulations of genetic data. The history of maize cultivation along with the areas and dates of first observations of WCR in North America were used to define 6 competing colonization scenarios differing in the combination of three main characteristics. First, the geographical origin of the species: WCR either originated in or near Mexico and expanded northward (“Mexican origin”), or it originated near present-day Colorado and expanded southward and eastward (“central-USA origin”). Because of the reduced number of samples in the southernmost area of WCR’s range, there is a risk that the true source population was not specifically sampled. Therefore, for all “Mexican origin” scenarios, we simulated sub-structuring within the oldest genetic unit as proposed by Lombaert *et al.* (2011). Second, the demographic history of the scenario’s first colonizing population: this population experienced either an “ancient bottleneck” (between 10,000 and 1,500 years BP) or a “recent bottleneck” (between 1,500 years BP and the date of first observation). This bottleneck could be the signal either of an introduction event from a native, unsampled, population or of a sudden decrease in population size during a selective sweep due to host plant shift. Third, the dates of the colonization events: either WCR accompanied the North American expansion of maize (“ancient expansion”, between 10,000 years BP and 1,500 years BP), or its range expanded only recently (“recent expansion”, between 1,500 years BP and the date of first observation). The competing scenarios thus differ in the direction of the colonization (south to north, or north to south) and by the relative recency of demographic and divergence events. In all scenarios, an expansion event corresponds to a simple divergence event from a source population possibly followed by a period at low effective size (bottleneck event) predating demographic stabilization at a higher effective size. Because the various populations under scrutiny are not separated by insurmountable geographical barriers, and because of the strong dispersal capacity of WCR (Coats et al. 1986; Grant and Seevers 1989; Bermond et al. 2013), we allowed continuous unsymmetrical migration between populations. All 6 scenarios are described in Table 1 and Figure S1.

**Table 1:**
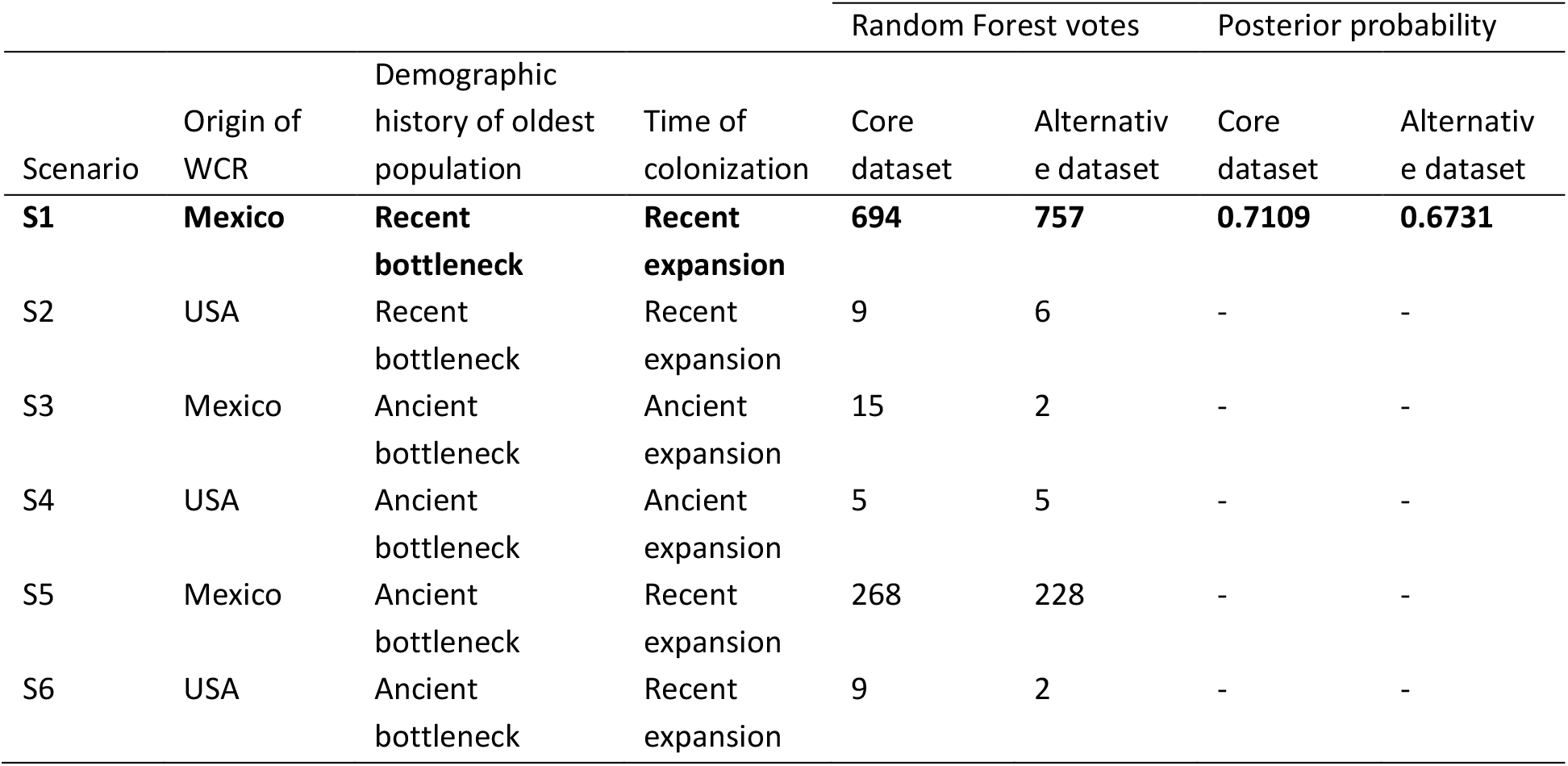
Description of the competing scenarios and results of the ABC analyses to infer the colonization history of WCR. Results are provided for both core and alternative datasets. The line in bold characters corresponds to the selected (most likely) scenario.

In our ABC analysis, historical, demographic and mutational parameter values for simulations were drawn from prior distributions defined from historical data and from a previous study (Miller et al. 2005), as described in Table S2. We used a total of 49 summary statistics: for each population (i.e. site-sample in the case of the observed dataset), we computed the mean number of alleles per locus, the mean expected heterozygosity (Nei 1987), the mean number of private alleles per locus and the mean ratio of the number of alleles to the range of allele sizes (Garza and Williamson 2001). For each pair of populations, we computed the pairwise *F*_ST_ values (Weir and Cockerham 1984) and the mean likelihoods of individuals from population *i* being assigned to population *j* (Rannala and Mountain 1997). For each trio of populations we computed the maximum likelihood estimate of admixture proportion (`Choisy et al. 2004). For all populations taken together, we computed the mean number of alleles per locus, the mean expected heterozygosity and the mean number of shared alleles per locus. These statistics were complemented with the five axes obtained from a linear discriminant analysis on summary statistics (Estoup et al. 2012).

To compare the scenarios, we used a random forest process (Breiman 2001) as described by Pudlo *et al.* (2016). Random forest is a machine-learning algorithm which circumvents curse of dimensionality problems and some problems linked to the choice of summary statistics (e.g. correlations between statistics). This non-parametric classification algorithm uses hundreds of bootstrapped decision trees (creating the so-called forest) to perform classification using a set of predictor variables, here the summary statistics. Some simulations are not used in tree building at each bootstrap (i.e. the out-of-bag simulations) and can thus be used to compute the “prior error rate”, which provides a direct method for cross-validation. Random forest (i) has large discriminative power, (ii) is robust to the choice and number of summary statistics and (iii) is able to learn from a relatively small reference table hence allowing a drastic reduction of computational effort. See Fraimout et al. (2017) and Momigliano et al. (2017) for recent case studies. We simulated 50,000 microsatellite datasets for each competing scenario, and checked whether the scenarios and priors were off target or not by comparing distributions of simulated summary statistics with the value of the observed dataset. We then grew a classification forest of 1,000 trees based on all simulated datasets. The random forest computation applied to the observed dataset provides a classification vote which represents the number of times a model is selected among the 1,000 decision trees. The scenario with the highest classification vote was selected as the most likely scenario. We then estimated its posterior probability by way of a second random forest procedure of 1,000 trees as described by Pudlo *et al.* (2016). To evaluate the global performance of our ABC scenario choice, we (i) computed the *prior error rate* based on the available *out-of-bag* simulations, and (ii) conducted the scenario selection analysis a second time with another set of site-samples (the “alternative dataset”) representative of the same genetic units as the core dataset, as suggested by Lombaert et al. (2014). Finally, we inferred posterior distribution values of all parameters, and some relevant composite parameters, of the selected scenario under a regression by random forest methodology (Raynal et al. 2017), with classification forests of 1,000 trees.

We used ABCsampler (Wegmann et al. 2010) coupled with fastsimcoal2 (v2.5, Excoffier et al. 2013) for simulating datasets and generating reference tables. We used Arlequin 3.5 (using the arlsumstat console version, Excoffier and Lischer 2010), in-house codes (perl and C++) and an R script used by Benazzo *et al.* (2015) to compute summary statistics. Scenario comparisons and parameter estimations were performed under R (R Development Core Team 2015) with the *“abcrf* package (v1.5, Pudlo et al. 2016).

Finally, as a control, we performed another ABC analysis with the same six scenarios using the software DIYABC (v2.1.0, Cornuet et al. 2014). In this context, simulations were run with no migration between populations, and the posterior probabilities of scenarios were estimated by polychotomous logistic regression (Cornuet et al. 2008) modified following Estoup *et al.* (2012).

## Results

### Genetic variation in WCR

The complete dataset, including a total of 917 individuals from 21 site-samples, displayed substantial polymorphism, with a mean of 12.69 alleles per locus, over all samples. Allelic richness corrected for 10 individuals ranged from 4.4 alleles per locus in a sample from Minnesota (MN) to 6.35 in a Mexican sample (MX-2). Overall, the southernmost site-samples displayed the highest diversities, especially in Mexico, and to a lesser extent in Arizona, New Mexico and Texas. Null allele frequencies were low with a mean of 0.017 for all locus-by-site-sample combinations. However, they were above 0.15 for two loci in the two Mexican site-samples, which very likely explain the larger F_is_ and significant Hardy-Weinberg tests. See Table S1 for a concise presentation of diversity measurements for each site-sample.

Genotypic differentiation was statistically significant in 137 of 210 pairwise comparisons between site-samples (Table S3). Global levels of differentiation between site-samples were moderate, with a mean F_st_ of 0.035. As previously described in other studies using lower numbers of samples and genetic markers (Kim and Sappington 2005; Ciosi et al. 2008; Kim et al. 2008; Coates et al. 2009), a large part of the northern USA, i.e. all site-samples north of the states of New Mexico and Texas, displayed high genetic similarity with a mean pairwise F_st_ of 0.005. In contrast, F_st_ values increased steeply with latitude, with the highest value (0.16) between site-samples MX-2 in Mexico and Mo-02 in Illinois (Table S3).

In the unrooted NJ tree, the position of the site-samples was mostly consistent with a latitudinal pattern (Fig. 2). Despite long branches, both Mexican samples grouped together, and were closest to Arizona, followed by New Mexico and Texas. The remaining 16 site-samples grouped together in a tight cluster with short branches. This pattern was supported by the significant correlation between pairwise genetic differentiation and geographic distance (P < 10^-4^).

**Figure 2:**
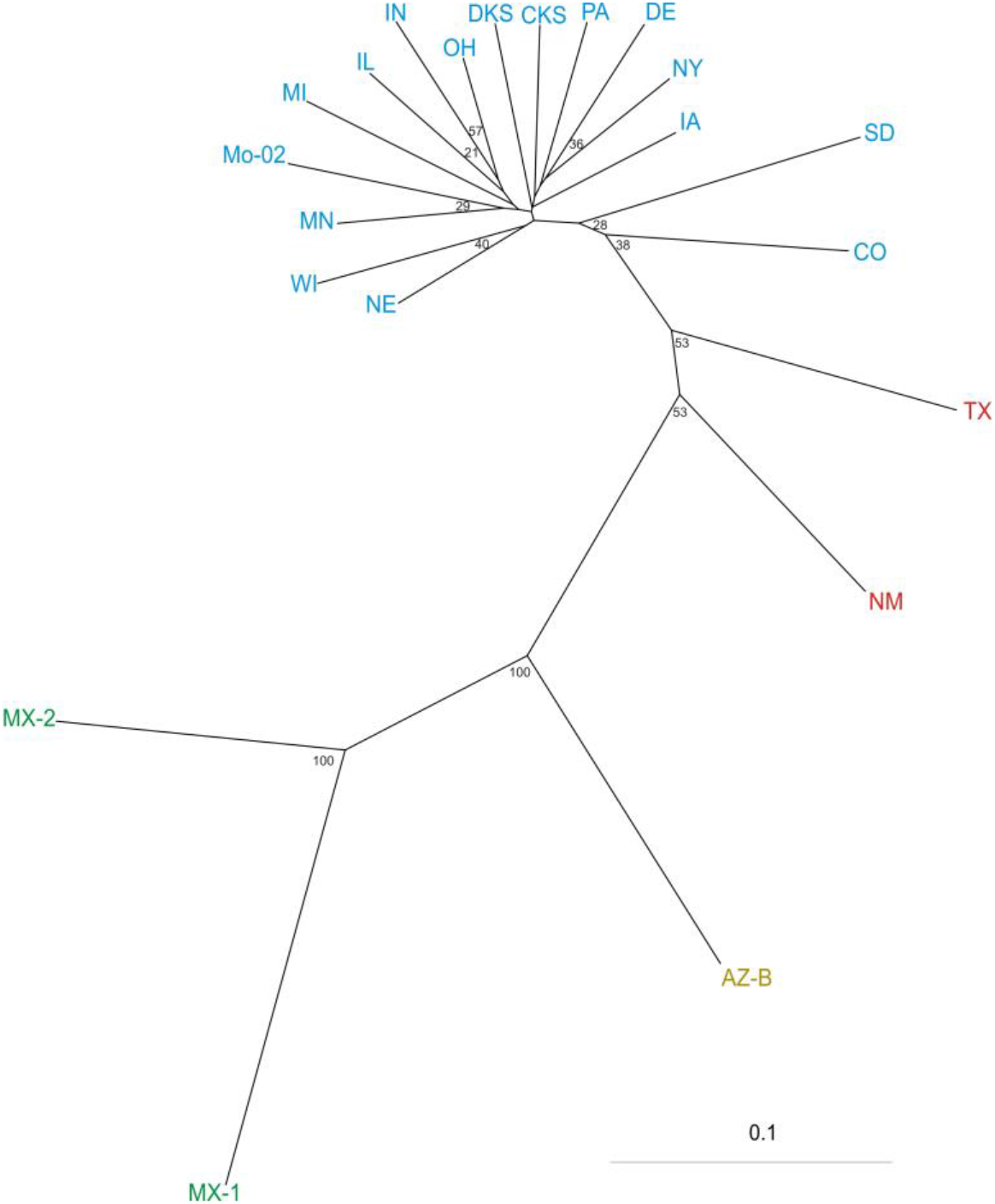
Neighbour-joining tree for WCR site-samples based on the chord distance of Cavalli-Sforza & Edwards (1967). Site-sample names are as in Figure 1 and Table S1. Site-samples of the same color belong to the same genetic unit as inferred from STRUCTURE and BAPS (Figures S3 and S4). Bootstrap values calculated over 1000 replications are given as percentages (only values >20% are shown).

### Population structure of WCR in North America

A hierarchical approach applied to both STRUCTURE (Pritchard et al. 2000) and BAPS (Corander et al. 2003) Bayesian clustering methods provided the same qualitative results. In the first round, site-samples were partitioned into three groups: the first contained MX-1, MX-2 and AZ-B site-samples, the second contained the NM and TX site-samples, and the third contained all 16 remaining site-samples. This partitioning is also observed at higher values of *K* (Fig. S2). Second rounds within each of these three groups only separated the two Mexican site samples (MX-1 and MX-2) from Arizona’s single site-sample (AZ-B). A third round showed no additional partitioning. Details of BAPS and STRUCTURE results can be found in Figures S3 and S4. To summarize, our 21 site-samples could be partitioned into four main genetic units clearly linked to geographical patterns (Fig. 1): (i) the “Mexico” genetic unit (46 individuals from 2 site-samples: MX-1 and MX-2), (ii) the “Arizona” genetic unit (40 individuals from 1 site-sample: AZ-B), (iii) the “New Mexico/Texas” genetic unit (82 individuals from 2 site-samples: NM and TX) and (iv) the “Colorado/New York” genetic unit (749 individuals from 16 site-samples: CO, DKS, CKS, NE, SD, IA, MN, WI, Mo-02, IL, IN, MI, OH, PA, DE and NY).

### Colonization history of WCR in North America inferred from ABC analyses

For the core dataset used in the ABC analyses, the choice of site-samples was based on the largest sample sizes for the “Mexico” and “New Mexico/Texas” genetic units: MX-2 and TX respectively. For the “Colorado/New York” genetic unit, we chose the site-sample CO from Colorado, because of its geographical proximity to the historical first observation of the species, and because of the well-described colonization history of this genetic unit eastward from this area (Gray et al. 2009). For the alternative dataset, the “Mexico” and the “New Mexico/Texas” genetic units were represented by the MX-1 and NM site-samples respectively, and the “Colorado/New York” genetic unit was represented by the OH site sample which displayed the lowest mean intra-genetic unit pairwise F_st_ (Table S3). In both datasets, the “Arizona” genetic unit was represented by the single AZ-B site-sample. Regarding the clear geographical partition of the four genetic units (Fig. 1), and the patterns observed in the NJ tree (Fig. 2), the “Mexican origin” scenarios represent a simple South to North expansion in this specific order: (i) “Mexico”, (ii) “Arizona”, (iii) “New Mexico/Texas” and (iv) “Colorado/New York”. The “central-USA origin” scenarios entail an expansion in the opposite direction, from North to South (Fig. S1). Raw dates of first observation were used as lower bounds of time prior distributions (Table S2): 1893 for “Mexico”, “Arizona” and “New Mexico/Texas” (i.e. 113 generations backward in time, Horn 1893), and 1867 for “Colorado/New York” (i.e. 139 generations back in time, Le Conte 1868). Depending on the topology of the scenario, these dates were narrowed by conditions.

Comparisons of distribution of simulated summary statistics with values of the observed core dataset showed that the combination of scenarios and prior that we chose was realistic: among the six simulated scenarios, we had from zero (scenarios 1 and 5) to only two (scenarios 2, 4 and 6) observed statistics out of 49 that significantly (at a 5% threshold) lay in the tails of the probability distribution of statistics calculated from prior simulations (Table S4).

The results of the random forest ABC analyses are shown in Table 1, and the selected scenario is graphically summarized in Figure 3. The results indicate, with a high probability of 0.71 for scenario 1, that (i) Mexico is the most likely first identifiable source of the colonization, (ii) a bottleneck occurred recently in this population and (iii) the colonization of North America by WCR is recent. The prior error rate was high (47.8%), but the result was qualitatively and quantitatively confirmed by the analysis of the alternative dataset which selected the same scenario with a very similar posterior probability (Table 1). This high prior error rate was caused by some scenarios being differentiated only by the prior distribution of divergence times. Indeed, the three “Mexican origin” scenarios (i.e. scenarios 1, 3 and 5; Fig. S1) brought together a total of 977 votes among the 1000 generated decision trees, with scenario 5 (i.e. ancient ancestral bottleneck and recent colonization) garnering the second highest number of votes. When comparing in a new analysis only the 3 scenarios with a Mexican origin differing by the times of colonization (scenarios 1, 3 and 5), scenario 1 with all historical events being recent obtained 743 votes among 1000. Finally, random forest ABC results were confirmed by the standard DIYABC analyses as well: scenario 1 was selected with probability of 0.935 and 0.939 for the core and alternate dataset respectively.

**Figure 3:**
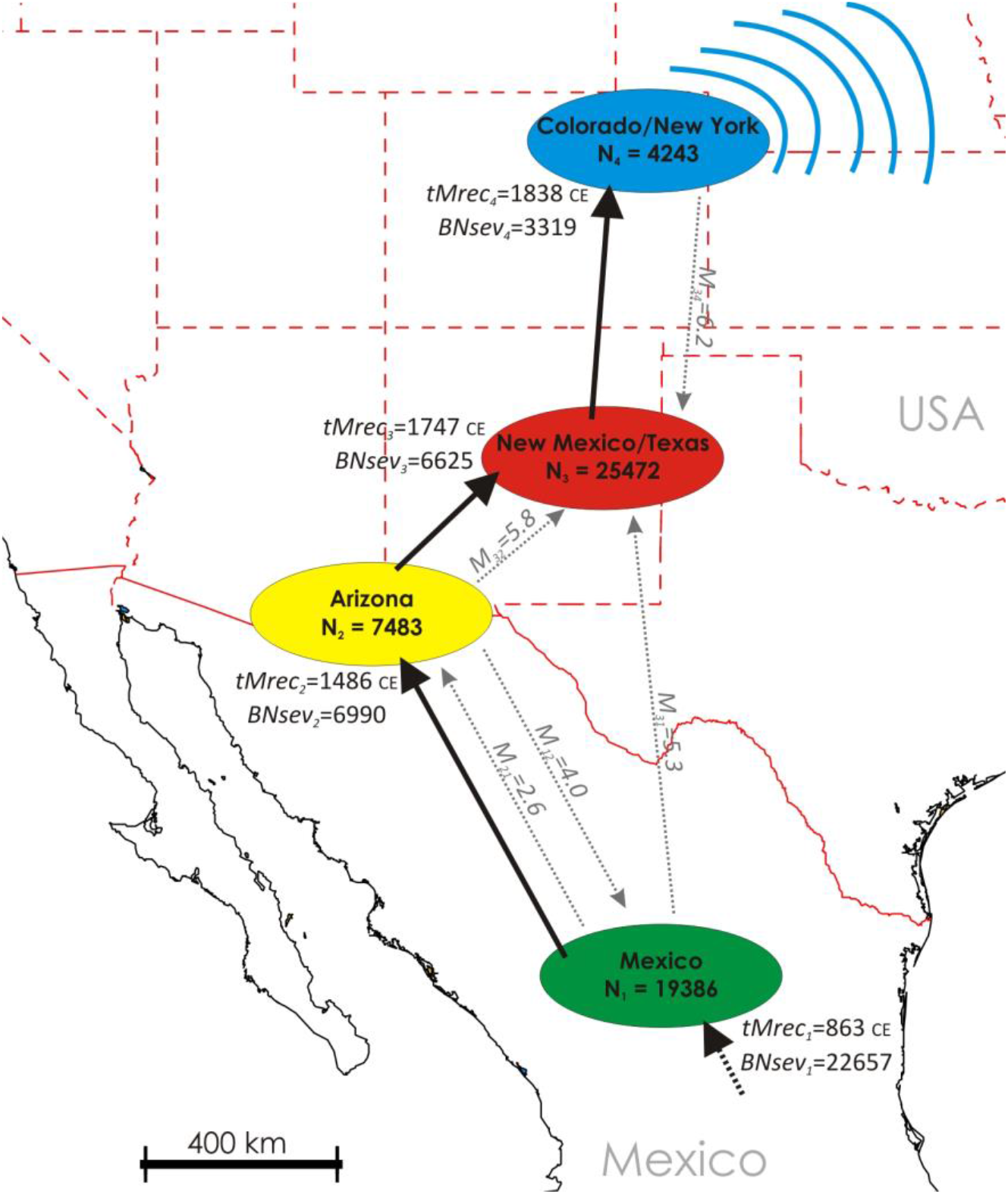
Graphical representation of the most likely scenario of WCR colonization of North America, and main parameter estimations. Notes: The four genetic units are those inferred from Bayesian clustering analyses. All parameter estimations were performed with samples MX-2, AZ-B, NM and CO representing the “Mexico”, “Arizona”, “New Mexico/Texas” and “Colorado/New York” genetic units, respectively. All displayed parameter values are the medians of posterior distributions (Table S5). *BNsev¡* = bottleneck severity of population *i* computed as *[BD_i_* × *N_parental population of population i_* / *NF_i_*]. *M_ij_* is the effective number of migrants per generation from population *i* to population *j* backward in time, computed as *m_ij_* × *N_i_*; only values above 2 individuals per generation are presented. All arrows are presented forward in time for ease of reading. Dates are presented in years of the Common Era (i.e. CE). Blue lines near the “Colorado/New York” genetic unit represent the well described eastward expansion after the 1940s (Gray et al. 2009).

Point estimates of key parameters from scenario 1 are presented in Figure 3 (complete results in Table S5). The “Mexico” genetic unit suffered a strong initial bottleneck probably around 1,100 years ago. The geographic expansion that followed northward was accompanied by successive bottlenecks of lesser severity than the ancestral one. Effective population size was lowest for the “Colorado/New York” genetic unit (median value of *N4* = 4,243 individuals) which is the more recent population. In contrast, the “New Mexico/Texas” genetic unit displayed the largest population size (median value of *N_3_* = 25,472 individuals). This geographically central population received the largest number of migrants from each of the three other genetic units (from 5.3 to 6.2 effective migrants per generation). Effective migration between genetic units was, however, globally moderate over North America (mean of all median effective number of migrants = 2.7 individuals per generation). Note that most parameter posterior distributions displayed large ranges (Table S5), so these results should be interpreted with caution.

## Discussion

The main results of our study are that the origin of WCR is in the south of its North American range, and that it has expanded northward. ABC results were indeed confirmed by those of more classical population genetics methods, such as the observation of a decrease in genetic variation from South to North, as expected from successive founder events during a range expansion (Le Corre and Kremer 1998; Hallatschek and Nelson 2008). This quantitative approach confirms what was previously proposed based on historical or phylogenetic data and rejects the hypothesis of a northern origin of WCR (Chiang 1973; Branson and Krysan 1981; Krysan and Smith 1987; Gray et al. 2009). However, our data do not allow us to determine the precise origin of the species. Our Mexican samples were collected in the state of Durango, while the WCR may have originated from further south in the country, or even in Guatemala. Indeed, the estimated strong ancestral bottleneck could be the signature of a first colonization step from an unsampled ancestral population.

Another important and unexpected conclusion of our study is that the history of WCR colonisation of North America is not associated with the early history of maize expansion out of Mexico into the American Southwest that began around 4,100 BP (Merrill et al. 2009; da Fonseca et al. 2015). Instead, our genetic data firmly indicate WCR did not arrive in the Southwest until about 1500 CE following an initial severe bottleneck detected in the Mexican sample at about 900 CE (Fig. 3). However, this time frame does strikingly correspond to the intensification of maize cultivation in the American Southwest, Great Plains, and Eastern Woodlands that began around 900 - 1000 CE (Fritz 1990; Boyd et al. 2008; Smith 2017). This widespread intensification of maize use was explosive (Simon 2017), and was probably related to the development of higher yielding varieties, which formed the basis of maize-dominated agricultural systems and more complex societies after 1000 CE (Smith 2017). Our analysis suggests that the most recent WCR population in the Colorado Great Plains region originated from colonization northward from New Mexico/Texas in the first half of the nineteenth century (Fig. 3). The absence of genetic structure that we observed from Colorado to New York is entirely consistent with the very recent colonization history by the species throughout this large area of great economic importance. This corroborates historical records (Chiang 1973; Metcalf 1983; Gray et al. 2009) and previous population genetics studies (Kim and Sappington 2005; Ciosi et al. 2008; Kim et al. 2008; Coates et al. 2009). It also explains the low estimated effective population size of the “Colorado/New York” genetic unit despite large population densities in the field, which is consistent with a still unmet mutation-drift equilibrium.

The reason for the seemingly late spread of WCR northward, thousands of years after maize was domesticated, is unclear. The genetic bottleneck suffered by the Mexico WCR population around 900 CE may be the signature of a very recent change of host from an unknown plant to maize. Alternatively, it may be a signal of expansion northward that may have depended on the more widespread planting of maize that began about 900 CE. The ability to grow nonrotated maize on the Great Plains was greatly enhanced in the mid-twentieth century by the introduction of sprinkler irrigation systems, soil insecticides, and synthetic fertilizers, and this triggered the eastward expansion of WCR (Gray et al. 2009; Meinke et al. 2009). Maize planted continuously in the same field (i.e., nonrotated maize) is a precondition for buildup of large populations of WCR (Branson and Krysan 1981; Levine and Oloumi-Sadeghi 1991), and thus large numbers of potential emigrants. A high proportion of nonrotated maize in the landscape also is important in facilitating establishment of an immigrant population (Youngman and Day 1993; Meinke et al. 2009). These circumstances created a habitat bridge that allowed the rapid eastward expansion of WCR into the rain-fed Corn Belt. The same principle, albeit over a much longer time scale, may have been at work in promoting the northward expansion of WCR out of Mexico when maize presence increased in the landscape post-900 CE.

In this paper, we have provided quantitative evidence for the first time of the southern origin of WCR in North America. Moreover, our results strongly suggest that the colonization of WCR in North America is very recent. Thus it appears that the species was not gradually co-domesticated with maize, but rather behaved as an invasive species. From its tropical origin, the species has quickly adapted to continental climates and has become one of the worst pests of maize. Considering the estimated chronology of the North American invasion, and the very likely underlying association with key modifications of maize cultural practices, WCR can be considered a product of modern agriculture, i.e. a recent man-made pest (Metcalf 1986).

## Acknowledgments

We thank our colleagues Rosanna Giordano, Stephan Toepfer, Uwe Stoltz, Kyung Seok Kim, Sue Ratcliff, Greg Cronholm, Lee French, Lance Meinke, Brendon Reardon, Eli Levine, Bruce Eisley, Dennis Calvin, Joanne Whalen and Ken Wise for *Diabrotica virgifera virgifera* beetles and DNA samples. We also thank Emeline Deleury, Arnaud Estoup and Andrea Benazzo for scripts to compute summary statistics. We also thank Alexandra Auguste, Paulette Flacchi and Marie-José Odonne for technical and administrative assistance. This work was funded by grants from ANR projects Bioinv4I and Emile, and from the French Agropolis Fondation (Labex Agro-Montpellier, BIOFIS).

## Data accessibility

Data associated with this article (including microsatellite data file, ABC reference tables, input and script files for performing ABC simulations and analyses) are archived in Zenodo:

http://doi.org/10.5281/zenodo.832120

## Author contributions

EL and TG designed the study. TS managed the collection of samples. MC, NM and AB genotyped the samples. EL and TG analysed the data. EL, MC, NM, TS and TG wrote the paper. All authors have revised and approved the final manuscript.

